# Immune System Alterations in Depression across Baseline and Flu Vaccine Challenge

**DOI:** 10.64898/2025.12.02.691874

**Authors:** Dhivya Arasappan, Alyssa Marron, Sina Sanei, Mbemba Jabbi

## Abstract

There are multiple reports of elevated inflammation in patients with major depression. It is, however, unclear whether these reported perturbations in immune functions in depression affect the general functionality of the peripheral immune system. Here, using single-cell RNAsequencing (scRNA-seq) of peripheral blood mononuclear cells (PBMCs) extracted from blood samples collected prior to a flu vaccination administration/immunization (at baseline), we found downregulated T cell and B-cell-associated gene expression repertoire coupled with a selectively upregulated plasmablast, CD14 classical monocytes, and natural killer (NK) proliferating cell-associated gene expression in 9 depressed patients (6 females) compared to 5 healthy controls (5 females), as well as in association with depression ratings. Our cell type proportion analysis revealed shifts in immune cell populations, specifically reduced numbers of CD4+ T and CD8+ T proliferating cells, plasmablasts, and NK cells in individuals with depression compared with controls. In contrast, we found increased numbers of CD14 classical and CD16 monocytes as well as doublet cells in depressed individuals compared with controls. Although our baseline and flu vaccine challenge did not show marked differences in peripheral immune markers measured by a multiplex cytokine assay, our results suggest impaired innate and adaptive immune responses at the transcriptomic and cellular population levels in depressed patients.

## INTRODUCTION

Depression is a prevalent and debilitating disease affecting about 350 million people globally at any given time (Herrman et al., 2022; Marwaha et al., 2023; Nemeroff, 2007). As one of the leading causes of disability-adjusted life years (DALYs) (Goldberg, 2011; Rush et al., 2006), depression results in an estimated annual economic burden of over 1 trillion dollars globally and is associated with ∼800 thousand global completed suicides annually (Xu et al., 2023). Although the prevalence of depression and the toll that depression inflicts on human health, socioeconomic output, and wellbeing, and ultimately suicide mortality is enormous, the pathophysiology of depression remains ill-defined.

Evidence of the involvement of inflammatory cytokines and other innate and adaptive immune mediators in depression is mounting (Kohler et al., 2017; Syed et al., 2018). In line with this emerging evidence of immunomodulatory perturbations in depression, marked increases in inflammatory markers have been observed in depressed individuals who have experienced childhood trauma or chronic stress (Andersen, 2022; Brown et al., 2021; O’Shields et al., 2023). However, the cumulative roles of the history of adverse childhood experiences like trauma and abuse, and the diagnosis of depression in adulthood (Smart et al., 2015) on specific inflammatory mechanisms remain unknown. Increased peripheral levels of C-reactive protein (CRP) (Ford & Erlinger, 2004) and both proinflammatory [e.g., interleukin (IL)-6 and tumor necrosis factor (TNF)] and are well-documented in depression (Beurel et al., 2020; Miller & Raison, 2023). At the cellular level, immunosuppression (Asnis & Miller, 1989; Beurel et al., 2020; Maes, 1995; Miller & Raison, 2023), as exemplified by decreased lymphoproliferative responses of T cells (Kronfol & House, 1989), lower Natural Killer cell activity (Irwin et al., 1990), and reduced numbers of T helper (Th) cells (Schleifer et al., 1989), has been reported in depressed patients. Yet, increased numbers of leukocytes (Kronfol & House, 1989), neutrophils (Irwin et al., 1990; Kronfol & House, 1989), and monocytes (Muller et al., 1989) have concurrently been documented in depression, suggesting a more complex picture across protein and cellular level measurements as pathophysiological markers for depression.

Both innate and adaptive immune mediators have been implicated in the pathophysiology of depression (Kohler et al., 2017; Miller & Raison, 2023; Syed et al., 2018). In particular, evidence of the role of IL-17A protein (Jha et al., 2017; Xie et al., 2023), which is produced by CD4 T cells and T helper 17 (Th17) cells (Chen et al., 2011) or other T cells (Kronfol & House, 1989), coupled with the role of cellular phenotypic and transcriptomic abnormalities, has been reported in depression (Bekhbat et al., 2022; Belzeaux et al., 2010). Notable findings in depression demonstrating its association with inflammation have been observational and cross-sectional (outside of the empirical framework of experimental manipulations of inflammatory processes) in approach, rendering the landscape of the inflammatory pathology in depression inconclusive. To uncover the potential immune dysfunctions associated with depression, we first confirm the existence of a Major Depressive Disorder (MDD) diagnosis or the absence of any mental health diagnosis in each participant using the Mini-International Neuropsychiatric Interview (MINI) questionnaire to identify depressed patients (N=17) and healthy controls (N=5). All participants were recruited from a community-dwelling volunteer sample. We examined the relationship between depression and a history of childhood adversity, as well as protein-level immune markers such as CRP and cytokines, by incorporating measures from the Childhood Adversity Questionnaire and protein-level assays in our group comparisons at baseline (i.e., prior to the influenza vaccination) and post-flu vaccine challenge. We then performed transcriptomic-level immune profiling of PBMCs using RNA sequencing on baseline samples collected from a sub-cohort of depressed patients (N = 9) and healthy controls (N = 5). We then sought to replicate the depression-related cell-type gene expression findings using baseline PBMC counts. We tested whether the immune response repertoire of depressed patients would be significantly perturbed relative to healthy controls at baseline and after administration of the inactivated influenza vaccine (IIV).

## METHODS

### Study overview

The study was conducted between January 2018 and November 2019 through the University of Texas at Austin (UTA) Mood and Anxiety Disorders Program (ClinicalTrials.gov ID NCT03756246). The UT Austin Institutional Review Board committee approved the study. All participants signed an informed consent form.

### Participants

#### Inclusion criteria

Immunologically healthy individuals with no prior history of immune/inflammatory conditions, ranging from 18 to 60 years old, who are eligible for but have not received the IIV within the past 12 months, were included in this preliminary clinical trial.

#### Exclusion criteria

All participants were screened at baseline and excluded if they had chronic or severe medical conditions, recent use of medications, or a history of smoking that could confound the results. All participants were asked to refrain from NSAIDs for 48 hours before baseline through the third day of participation. Participants with elevated white blood cell count or other indicators of recent infection or occult illness (e.g., elevated fasting blood glucose) were excluded from the follow-up repeated-measures study.

Of the 36 eligible participants who were recruited, 22 completed the baseline, IIV challenge, and follow-up components of the study. The sample consisted of 17 participants diagnosed with major depressive disorder (MDD) and 5 healthy controls (Table 1). Given the almost two-year stoppage of human research activities in 2020 and 2021, coupled with the potential population-level changes in people’s post-COVID-19 immune and inflammatory landscape, this study could not include additional participants beyond the 22 people who completed the study before the outbreak of the pandemic, as recruitment and data collection were permanently discontinued at the outbreak of the COVID-19 pandemic.

**Table 1.** Demographic characteristics of MDD vs. control samples that completed the baseline measures before receiving the flu vaccine.

|  | HC<br>n=5 | MDD<br>n=17 |
| --- | --- | --- |
| Participants | 5 | 17 |
| Gender (F/M) | 5/0 | 14/3 |
| Age | 22.2±1.3 | 30±3.7 |
| BMI | 23±1.3 | 26.5±1.3 |
| Race, n (%) |  |  |
| Caucasian | 40 (%) | 53 (%) |
| African-American | 20 (%) | 12 (%) |
| Asian | 20 (%) | 12 (%) |
| Other | 20 (%) | 23 (%) |
| Ethnicity |  |  |
| -Hispanic | 20 (%) | 23 (%) |
| -Non-Hispanic | 80 (%) | 77 (%) |
| Depression History |  |  |
| QIDS | - | 16.1±0.6 |

Complete blood count and other screening labs were obtained at baseline (Suppl Table 1). All participants underwent the same diagnostic interviews and IIV challenge. Participants completed the MINI diagnostic interview (Sheehan et al., 1998) and the Quick Inventory of Depression Symptomatology – Self Report (QIDS-SR) during the baseline visit. Among the patients with major depression, a score of 13-19 on the QIDS-SR was considered a current episode (a time of rating administration) of moderate depression, while a QIDS score of >=20 was regarded as a current experience of severe depression. Suicidal individuals were referred to specialty care (and excluded from the study) according to existing protocols. The Childhood Trauma Questionnaire (CTQ), a widely used 28-item selfreport questionnaire that assesses the history of experiences of abuse or neglect in childhood (Bernstein et al., 1994), was administered to all participants. It consists of five subscales measuring emotional, physical, or sexual abuse and emotional or physical neglect (Bernstein et al., 2003). It exhibits good internal consistency, and each subscale contains items assessing minimization and denial, with all CTQ items reported on a 5-point scale ranging from "Never True" to "Very Often True. "The scale requires approximately a 6^th^-grade reading level to complete. We used the total scores for the CTQ and the three abuse subscales measuring the history of emotional, physical, or sexual abuse for our analyses.

Additionally, we generated present/absent ratings based on the criteria established by Walker and colleagues (1999). These cutoffs include scores of 8 or more on sexual and physical abuse and 10 or more on emotional abuse (Walker et al. 1999). Healthy participants were screened to have no history of primary Axis I diagnosis as assessed with the MINI (Sheehan et al., 1998) and had CRP levels <3 mg/L at screening. All 22 participants of the influenza vaccine study received a single injection of the trivalent Afluria IIV challenge (Seqirus, Parkville, Australia) into the deltoid muscle at the end of the screening/baseline visit. Based on evidence in the literature, the inflammatory component of the response to the IIV challenge peaks one day after vaccination in healthy humans, as shown in both cytokine levels (Christian et al., 2015) and gene expression analysis (Bucasas et al., 2011; Nakaya et al., 2011; Tsang et al., 2014). A four-time point measurement was therefore adopted (baseline, followed by days 1, 3, and 28 blood sample collection) to capture the gradual changes in IIV-induced inflammatory responses reliably over the course of one month.

#### Blood Sample Collection, PBMC Isolation, and Storage

Blood samples were collected from the 22 participants between 8:30 and 10:30 am. Participants fasted for at least 8 hours before the laboratory visits and refrained from vigorous exercise for the preceding 24 hours. Blood was diluted 1:5 in Phosphate-buffered saline, underlaid with 10mL of Lymphoprep or Ficoll. The samples were centrifuged for 20 minutes at 1200 g without interruption at room temperature. The buffy layer of cells was recovered, washed with RPMI medium, and centrifuged for 10 minutes at 500 g at 14 °C. Red blood cells were lysed using ACK buffer for 5 minutes, and the PBMCs were filtered through a 40 μm filter. The cells were counted using a cell counter, aliquoted, and immediately stored at -80°C, then long-term stored in liquid nitrogen. Serum and plasma were aliquoted and stored at -80°C.

#### Cytokine/chemokine/growth factor analyses

Plasma cytokine, chemokine, and growth factor levels were measured on a MAGPIX (Luminex) using the human 27Plex cytokine multiplex assay (M500KCAF0Y, Bio-Rad) according to the manufacturer’s protocol. Assays were checked for quality control to fit the standard curve. Serum CRP concentrations were obtained by using the particle-enhanced immunoturbidimetric GEN 4 assay on a Roche Cobas analyzer by the Clinical Pathology Laboratory in Austin, Texas, USA.

#### Single-cell library preparation and cell-type RNA sequencing

Among participants who completed the repeated-measures study, including the IIV challenge (N = 22), we conducted single-cell PBMC RNA sequencing for only a subset (N = 14). PBMCs of 5 healthy controls and 9 depressed patients were selected from the 22 samples that participated in the IIV challenge study. To ensure an unbiased selection of the sub-sample of depressed patients for inclusion in the PBMC RNA-seq analysis, we selected 5 healthy controls and 5 MDD participants matched to the healthy controls by age. We then selected 4 additional MDD participants who most closely matched the age of the 5 healthy controls. This matching effort excluded all participants aged 27 or older (all of whom had MDD), as well as one 24-year-old participant with MDD.

To perform the single-cell RNA sequencing, PBMCs of the selected participants (5 controls and 9 MDD) were thawed and resuspended in RPMI 1640 media supplemented with 10% FBS, 100 IU/mL penicillin, 100 μg/mL streptomycin, 1 × non-essential amino acids, 1 μM sodium pyruvate, 2.5 μM β-mercaptoethanol, and 2 mM L-glutamine for 1 hour. Dead cells were eliminated using the Dead Cell Removal Kit (Miltenyi Biotec). Cell suspensions were assessed using the Nexcelom Cellometer K2. Viability ranged from 70.5% to 93.8% across samples (**Suppl. Table 1**). The suspensions were prepared following the 10X Genomics Chromium Next GEM Single Cell 5’ Reagent Kits v2 User Guide. First, cells were partitioned into nanoliter gel beads-in-emulsion (GEMs). Libraries were then created and sequenced on the Illumina Novaseq 6000, using 10X barcodes to assign reads to individual partitions and, consequently, to each cell. We generated 14 libraries, with ∼5,000 sequenced cells/sample at ∼50,000 reads/cell or ∼2,000 median genes/cell (**Suppl Table 1**).

### Statistical Analysis

#### PBMC Cell-Type RNA-Sequencing Data Analysis

Cell Ranger (Zheng et al., 2017) counts were generated for each Chromium FASTQ file. The pipeline mapped reads to the human (GRCH38) transcriptome using STAR (Dobin et al., 2013) and performed barcode counting and UMI counting to generate a counts matrix for each sample. Each count matrix was read as a Seurat object (Butler et al., 2018), and all the objects were combined using the Seurat merge function. SCTransform (Hafemeister & Satija, 2019) was used to normalize the cell-to-cell variation in the data, which could be due to technical confounds. We applied a modeling framework for the normalization and variance stabilization of molecular count data from singlecell RNA-seq experiments in line with previously validated methods (Hafemeister & Satija, 2019). The Seurat multimodal reference mapping pipeline (Hao et al., 2021) was used to identify cell-type clusters from the data. A PBMC dataset (Hao et al., 2021) released by 10X Genomics was used as the multimodal reference dataset. The reference dataset was also normalized using SCTransform, and anchors were identified between the reference and our dataset using the FindTransferAnchors function. The cell type labels were then transferred from the reference to our dataset using the MapQuery function, and labels were generated for our clusters at two levels of granularity (l1 and l2). Uniform Manifold Approximation and Projection (UMAP) was used for dimension reduction to visualize cell-type clusters. Top markers were identified for every cluster using the FindMarkers function. Markers between the diagnostic conditions (MDD vs healthy controls ‘HC’ groups) were also identified using the FindMarkers function and the DESeq2 (Love et al., 2014) method, and the Wilcoxon signed rank nonparametric test was used for group comparisons. Markers with log2 fold change between 1 and 20 and p-value less than 0.01 were identified as upregulated markers, and markers with log2 fold change between -1 and -20 and p-value less than 0.01 were identified as downregulated markers. Enrichr (Chen et al., 2013) was used for each marker set to identify enriched functional terms among the differentially expressed PBMC markers.

To determine whether measures of low-grade inflammation affected the overall immune system landscape in depression, we profiled baseline PBMCs by single-cell RNA sequencing using the 10X Genomics approach. Unsupervised clustering and UMAP projections were conducted using the 73,347 PBMCs from the 5-9 biological replicates of 14 participants in a single dataset (**Fig. 1A**). This analysis generated an average of 50,950 reads per cell and 2,154 genes per cell. Cells were sequenced to comparable depths and showed similar median molecular identifier counts and median gene numbers across all conditions. The origin of each cell type (**Fig. 1A**) was visualized in color-coded UMAP plots using the canonical markers of the different immune cells present in the blood.

**Figure 1:**
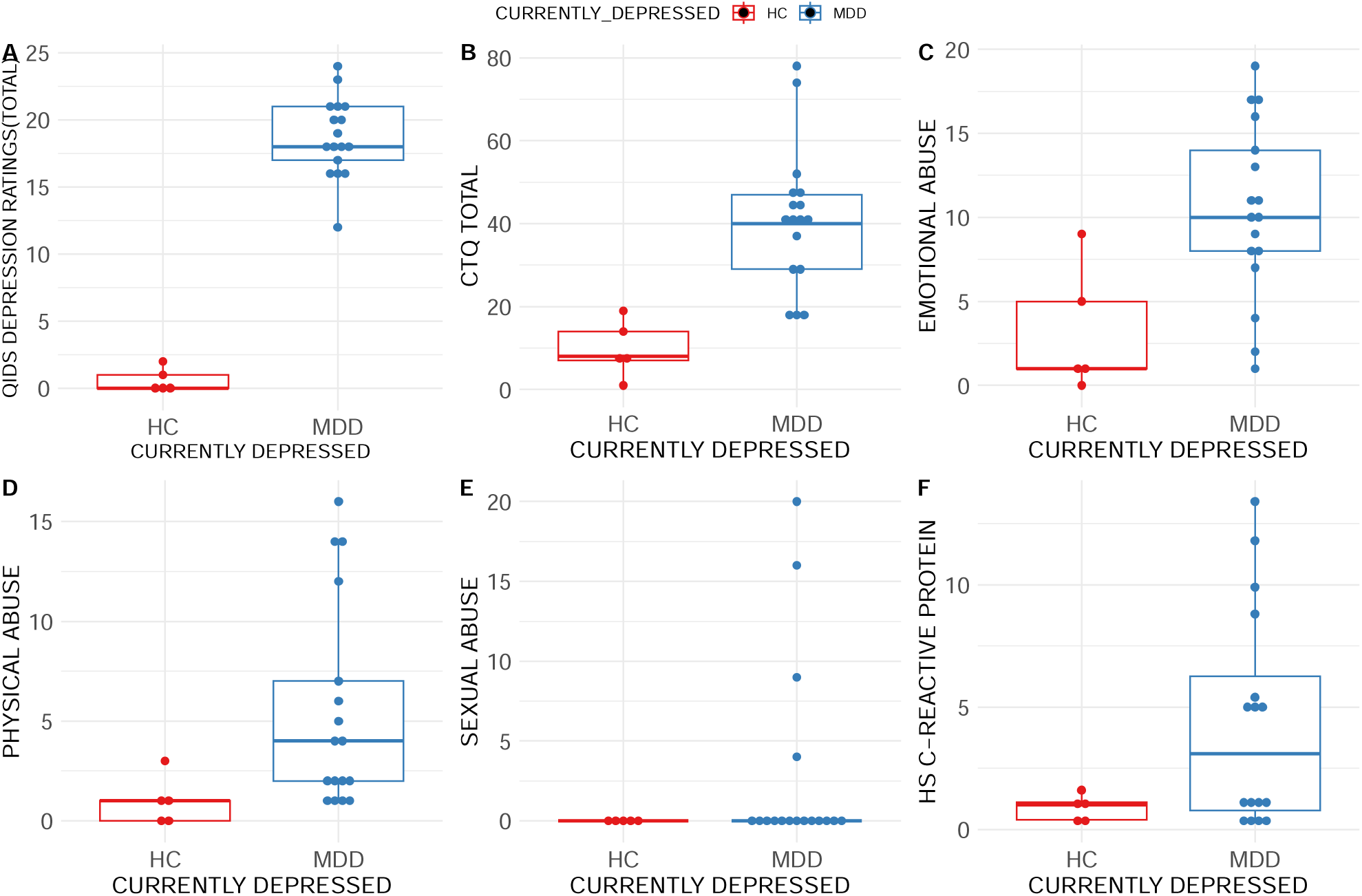
**A**, QIDS score; **B**, Childhood trauma history; **C**, Childhood emotional abuse history; **D**, Childhood physical abuse history; **E**, Childhood sexual abuse history; **F**, serum level of CRP, in depressed patients (MDD) vs. controls at baseline. Data were presented as means±SEM. n=5-17 participants/group, non-parametric permutation tests *p<0.05.

A permutation test was used to compare the proportions of each identified cell type between MDD and healthy controls, and cell types with an absolute log2 fold difference > 0.58 and FDR < 0.05 were identified as having significantly differential proportional counts in depressed patients compared to healthy controls. The single-cell RNA sequencing data will be deposited and made freely available through NIH data archives (NDA).

#### Data Transformation

For each variable, data were presented on a natural scale, and we adopted an approach that involves a ‘baseline measurements + 1’ transformation to address zero measurements in some variables. To reduce deviations from normality across all data types, we first applied the Box-Cox transformation using a grid search approach to identify the optimal power transformation parameter(λ) to baseline (before administering the IIV) data for all variables. For the Box-Cox transformation, we adopted the condition that, if λ was estimated to be zero, a log transformation was used; otherwise, the Box-Cox transformation was applied. Once the optimal λ was estimated, we transformed the baseline data and any available postimmunization repeated measurements. The box-cox() function from the MASS library in R version 4.3.2 was used for the transformation procedures (Ripley & Brian 2023).

#### Statistical Testing for Multiple Measurements

A linear model was used to test the significance of the intervention (IIV/time) across the post-vaccination days 1, 3, and 28. Participants were considered as random effects, and age was included as a covariate in the models to adjust for baseline age differences for all variables. The time effect was removed from the model for variables measured only at baseline. Among the 6 baseline-only measures, 4 (i.e., the quick depression inventory scale ‘QIDS’; composite childhood trauma questionnaire ratings ‘CTQ’; and the subscale of physical and emotional abuse experience history subcomponents of the CTQ) showed a significant condition effect. Out of the 28 repeatedly measured variables, only 3 demonstrated at least one significant main effect or interaction after permutation tests.

#### Permutation Tests

Permutation tests were performed using the permuco package in R, which offers various permutation methods (Kherad-Pajouh & Renaud, 2010). For baseline-only measures, the “freedman_lane” method was used, which is the default in the permuco package.

For repeated measures ANOVA, the “Rde_kheradPajouh_renaud” method was employed.

## RESULTS

### Depression diagnosis is associated with early life adversity and inflammation

Our permutation tests confirm that depressed individuals reported more symptoms than controls, p = 0.0002; F = 123 _(df1, 19)_ (**Fig. 1A**), as measured with the QIDS scale, which is an obvious result. Further assessment of the history of experienced childhood trauma as measured with the CTQ questionnaire revealed increased overall childhood trauma experience in the depressed patients vs. healthy controls, p=0.0004, F=18_(df1,19)_ (**Fig. 1B**). Emotional abuse history, p=0.015, F=7.4_(df1,19)_ (**Fig. 1C**). Physical abuse history, p=0.017, F=6.78_(df1,19)_ (**Fig 1D**) both were found to be higher in depressed patients. In contrast, sexual abuse history was not reported to be significantly different between depressed patients and healthy controls in our studied cohorts, p=0.5, F=0.47_(df1,19)_ (**Fig. 1E**). Of note, we found no difference in CRP levels (**Fig. 1F**) or cytokine levels (**Supplementary Fig. 1**) between depressed patients and healthy controls when controlling for age before the administration of the IIV/baseline or post challenge.

### Differentially expressed genes in populations of PBMCs affected in depression at baseline

Using the Wilcoxon parametric test to identify differential expression between conditions in R, we found that specific genes such as NFKBIA, DUSP1, GPX4, PLIND2, and ZFAND5 (and other genes in the top 50 downregulated genes) as well as predominantly more downregulated FKBP11, and ADAM8 genes in CD14 classical monocytes (**Fig. 2C**) in the depressed versus (vs.) controls. We further observed predominantly more upregulated set of genes such as RNADBP3, and YAF2 etc. in plasmablasts cells in depressed vs. controls (**Fig. 2E**). Using the individual scores of the quick depression inventory scale (QIDs) to estimate symptom severity at baseline as regressors, we identified a diverse set of genes such as NME2, and RBM3 etc. that were dysregulated in individuals with a depressive disorder relative to controls, in innate immune CD14 and CD16 monocytes, NK cells, and B memory cells, as well as in adaptive immune CD4 and CD8 proliferating (Schleifer et al. 1989) and plasmablast cell types (see **Fig 2G** for the cell-type specific gene expression profiles) were driven by the degree of self-reported depressive symptom severity in the total sample.

**Figure 2:**
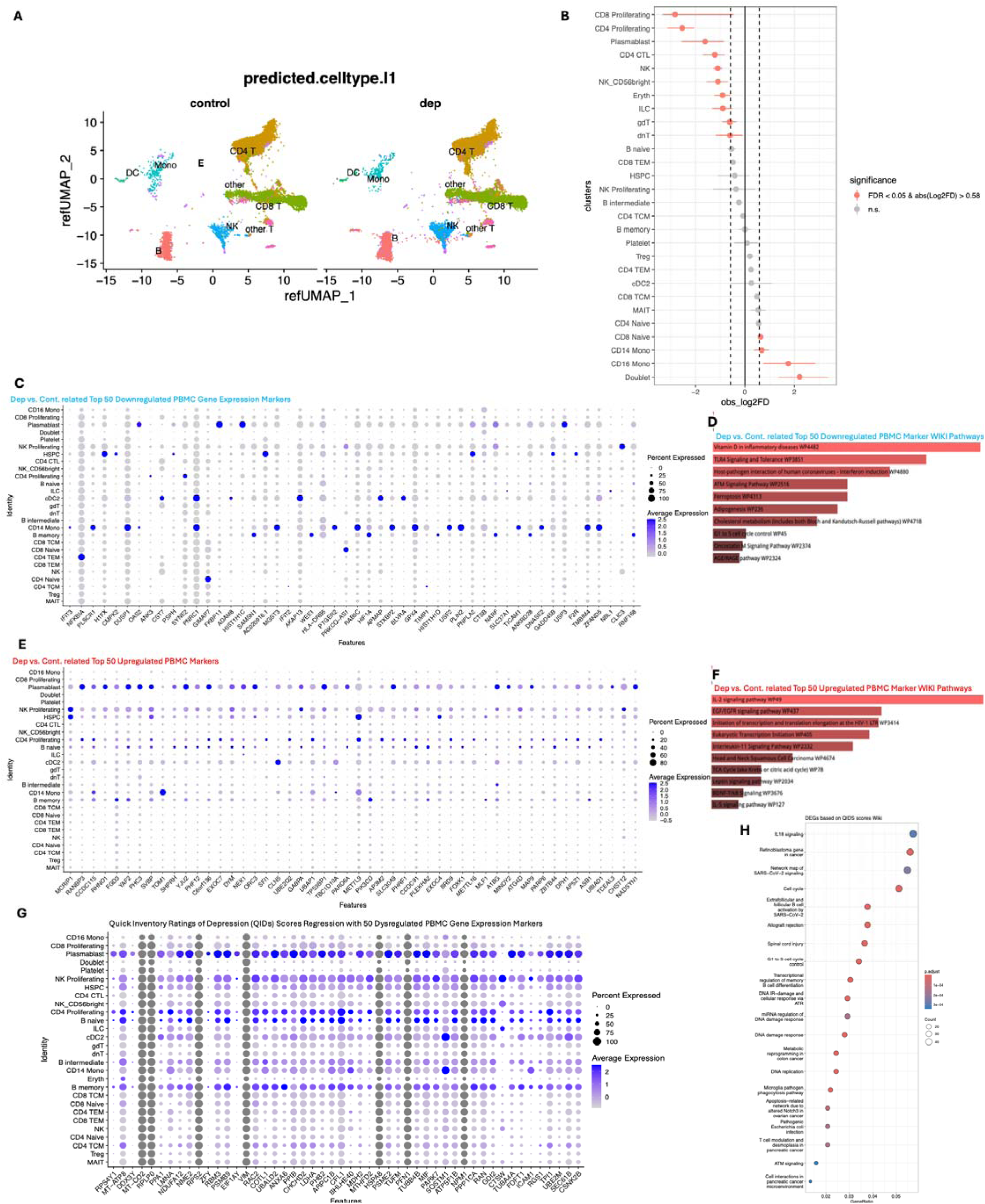
Single-cell RNA sequencing analysis of PBMCs of depressed patients and healthy controls. Single-cell RNA sequencing of PBMCs in 9 depressed patients (MDD) and 5 healthy controls (HC) was performed using the 10X Genomics sequencing. **A**, UMAP projections of single-cell RNA-Seq data, colored by cluster and labeled by cell type, separated by condition. **B**, PBMC cell type population frequencies indicated in depression vs. control revealed that CD4 T and CD8 T proliferating cells, plasmablasts, and CD4 CTL, as well as natural killer (NK) cells, were proportionally reduced in cell count in depressed participants. In contrast, CD14 (to a lesser extent) and CD16 (predominantly) monocytes were increased in cell count in the depressed, with the FDR and log2 fold change statistical thresholds used to compare cell-type proportional counts in the depressed compared to controls (significant differences indicated by red dots). **C**, Depression vs. controls comparison showing the top 50 PBMC RNA-seq markers downregulated in depression, especially in CD14 classical monocytes (also increased in cell count) and B memory cells. **D**, WIKI pathways representing the top 50 downregulated PBMC markers in depression, including reduced ATM signaling and G1 to S cell cycle control. **E.** Depression vs. controls comparison showing the top 50 PBMC RNA-seq markers upregulated in depression, reflecting significant upregulation of genes in adaptive immune plasmablasts (also decreased in cell count) and in innate immune NK proliferating cells (also reduced in cell count) in depressed participants. **F**, WIKI pathways representing the top 50 upregulated PBMC markers in depression, including transcription and translation elongation initiation, IL2, IL-5, and IL-11 signaling, as well as BDNF-TrkB signaling. **G**, Using the individual scores of the Quick Depression Inventory Scale (QIDS) to estimate symptom severity (baseline scores) as regressors, the results identify the top 50 genes whose differential expression across PBMC cell types was driven by individual differences in self-reported depression severity. **H**, WIKI pathways representing the top 50 dysregulated PBMC cell type-associated gene expression markers showing an association with increased self-reported QIDs scores of depression include ATM signaling and G1 to S cell cycle control pathways.

#### PBMC Single Cell Pathway Analysis

We then performed pathway enrichment analyses of the observed PBMC differential gene expression related to depression vs. controls, and QIDs association, and found that our observed downregulated PBMC markers in depression (**Fig. 2C**) were related to reduced ATM signaling pathways and G1 to S cell cycle control pathways (**Fig. 2D**). Further analyses of the WIKI pathways representing the upregulated PBMC gene expression markers observed in depression included transcription and translation elongation initiation pathways, IL2, IL-5, and IL-11 signaling pathways, as well as BDNF-TrkB signaling (**Fig 2F**). Finally, our analysis using WIKIpathways identified the observed dysregulated PBMC gene expression markers associated with increased QID scores for depressive symptoms as being related to ATM signaling, G1-to-S cell cycle control signaling, and several immunometabolic pathways (**Fig. 2H**).

### Depressed patients exhibit proportionally increased CD14 and CD16 monocyte counts, whereas CD4 and CD8 T cells, plasmablasts, and NK cells are proportionally decreased compared to controls

Applying PBMC single-cell RNA-sequencing measures of gene expression using the cell-count-proportional test in R, we identified and compared immune cell populations between depressed patients and healthy controls. We found increased proportional representation of CD14 and CD16 monocytes, as well as naïve CD8 T cells (all surviving p < 0.05 FDR at log2 fold change, 0.58) in depressed vs. controls, consistent with the findings of a recent study showing a spike in monocyte cell counts in depressed individuals (Bekhbat et al., 2022). In contrast, the populations of CD8 proliferating T cells, CD4 proliferating T cells, plasmablasts, central memory (TCM) CD4 CTL cells among CD4 cells, NK cells, unconventional T cells (dnT), gamma-delta T cells (γδT), and innate lymphoid cells (ILC) were decreased in counts in depressed patients compared to healthy controls [p<0.05 false discovery rate corrected ‘FDR,’ at log2 fold change (0.58)] (**Fig 2B**).

These findings confirmed the immunosuppressive phenotype previously reported in depression (Irwin et al., 1990), with a decrease number of proliferating T cells in depression and the associated reduction of T cell memory formation (Maes, 1995; Syed et al., 2018) coupled with an increase of monocytes consistent with the hyperactivation of the innate immune system and a possible defect in T cell priming.

## DISCUSSION

In the present study, we investigated the immune landscape before and after IIV administration in patients with depression and compared it with that of healthy controls. At baseline, just prior to IIV administration, depressed patients reported significantly more depressive symptoms and higher self-reported experience of childhood emotional, physical, and composite trauma history than healthy controls. These observed increments in self-reported levels of both depressive symptoms and childhood trauma history in the depressed patients compared to healthy controls replicated previous results (Souama et al., 2023; Wang et al., 2023) and supported the hypothesis that prior childhood trauma is a likely risk factor for later life depressive disorders (Teicher et al., 2022; Williams et al., 2016). Baseline CRP levels, a marker of low-grade inflammation, were not elevated in depressed patients compared with healthy controls after correction for age, confirming that CRP concentration increases with age (Lowe, 2005). In line with our CRP results not being elevated in the depressed sample when age is corrected for, other protein-level cytokine markers were not elevated in depression, both at baseline and in response to the flu vaccine over time. The duration of depression and the number of depression episodes, which are all likely age-dependent, also influence the levels of CRP (Karlović et al. 2012) and other inflammatory markers. Regarding cytokine production, it is possible that the inflammatory cascade has not yet been initiated in these depressed patients, and the increased cytokine production may result from an immune cell defect. Indeed, when analyzing the immune cells from the blood of depressed patients and healthy controls, we found baseline immune cell type gene expression dysregulation in depressed patients, including increased proportional count of innate immune classical CD14 monocytes, and non-classical CD16 monocytes, coupled with a decreased proportional count of various adaptive immune cells (e.g., CD4 proliferating, CD8 proliferating cells, plasmablasts and NK cells) at baseline.

At baseline, we identified differentially expressed genes in depressed patients compared to healthy controls, including NFKBIA (a transcription factor that regulates immune/inflammatory genes), DUSP1, GPX4, PLIND2, and ZFAND5, and more downregulated FKBP11 and ADAM8 genes in CD14 classical monocytes. In contrast, we observed a more upregulated gene set, including RANDBP3 and YAF2 (among others), in plasmablast cells from depressed individuals relative to controls. These cell-type-specific gene expression results, at least at the pathway enrichment level, were driven by individual scores on the Quick Depression Inventory Scale (QIDS), implicating immune and inflammatory pathway abnormalities. Of interest, individual differences in self-reported scores of depressive symptoms (as measured with the QIDs scale) predicted a diverse set of PBMC gene expression dysregulations including innate immune CD14 classical monocytes and CD16 monocytes, NK proliferating, and B memory cells, and cells, as well as dysregulated adaptive immune CD4 and CD8 proliferating (Schleifer et al. 1989), and plasmablasts cellular gene expression. Akin to depression-related neuroinflammation, previous reports showed that the CD14 and CD16 monocyte phenotypes in depression were associated with microglial pathogen phagocytosis pathways, suggesting a monocyte-linked proinflammatory property in depressed patients (Rossol et al., 2011; Segura et al., 2013).

Recapitulating our PBMC gene expression analysis, we found that depressed patients had fewer CD4 and CD8 proliferating T cells than controls. In contrast, depressed patients showed increased numbers of monocytes relative to healthy controls. Recent studies are increasingly pointing to elevated monocyte counts in association with both psychosocial stress experience (Wohleb et al. 2015; Ramirez et al. 2017; Hu et al. 2022; Cathomas et al. 2024), fatigue (Berentschot et al. 2023), depression (Muller et al., 1989; Foley et al. 2023), and in depression-related phenotypes, including increased suicide risk (Puangsri & Ninla-Aesong 2021). In line with our convergent findings of monocyte gene expression changes in depression, coupled with a nearly two-fold increase in monocyte count, we also observed convergent changes in T-cellular measurements across different levels of depression. Specifically, we found altered CD4 T and CD8 T cellular gene expression across the transcriptome, as well as elevated CD4 T and CD8 T cellular immune responsivity measures, and over two-fold decrements in both CD4 T proliferating and CD8 T proliferating cell counts in depression. Our findings of convergent T cellular and monocyte related abnormalities in terms of proportional count decrements of T cellular counts and increments in proportional monocyte count in depression, are in line with earlier findings of monocyte abnormalities that are linked to observed peripheral monocyte trafficking/infiltration of key brain regions (Wohleb et al. 2015; Ramirez et al. 2017; Hu et al. 2022; Cathomas et al. 2024), and together points to multi-level monocyte abnormalities as fundamental pathobiological mechanism for depression. Together, these findings are consistent with the thesis that immune/inflammatory mechanisms are dysfunctional in some depressed individuals across different measurement scales (Bekhbat et al., 2022).

Our findings of an increased proportional population of monocytes in the blood of depressed patients replicate previous studies (Bebhkat et al., 2022). Consistent with recent studies, elevated monocyte counts are associated with psychosocial stress experience (Wohleb et al., 2015; Ramirez et al., 2017; Hu et al. 2023; Cathomas et al., 2024), fatigue (Berentschot et al., 2023), measures of depression (Muller et al., 1989; Foley et al., 2023), and specific depression-related phenotypes like increased suicide risk (Puangsri & Ninla-Aesong, 2021). Our findings of proportional increments and gene expression changes of monocytes in depression vs. controls are strongly in line with earlier findings of monocyte abnormalities leading to monocyte trafficking/infiltration of key brain regions (Wohleb et al. 2015; Ramirez et al. 2017; Hu et al. 2023; Cathomas et al. 2024), and point to multi-level monocyte abnormalities as a potential fundamental pathobiological mechanism in depression. Together, these findings are consistent with dysfunctional immunity/inflammation in depressed individuals (Bekbhat et al., 2022).

The present study has several strengths and limitations. While only our protein-level measurements were analyzed for baseline and post-flu vaccine challenge, these measurements did not reveal significant depression-related inflammatory effects that survived multiple corrections. Whether the small sample size could have deflated our lack of protein-level immune deregulatory findings needs to be assessed with larger samples. Our intensive characterization of the immune the landscape at the transcriptomic, protein, and cellular phenotypic levels before and after the IIV challenge in depressed individuals and healthy comparison samples did yield a novel insights related to single-cell transcriptomics revealing novel cell-type CD4 T cell CD8 T cell, plasmablasts, as well as monocyte gene expression dysregulations in depressed individuals coupled with proportional changes in cell type proportional counts in the depressed samples but only at baseline. In summary, we report in this proof-of-concept study that depressed patients, compared to healthy controls, exhibit more RNA-level and cell-population-level immune and inflammatory CD4 T-cell, CD8 T-cell, plasmablast, and monocyte repertoires associated with depression at baseline. These findings revealed a complex depression-related multicellular immune mechanism at the cell phenotypic and transcriptomic levels than previously thought.

## Acknowledgments

This trial was sponsored by the NIMH (Grant #R21MH115326). We thank Roy D Mayfield for supporting this project’s data storage and analytic resources. We sincerely thank our colleagues at the University of Miami sequencing facility for performing the single-cell RNA sequencing and the Multiplex assays.

## Author contribution

DA, MJ, SS, and Paul Rathouz performed the data analysis. MJ, AM, and DA wrote the manuscript with inputs from SS and Paul Rathouz. AM and Dr. Marisa Toups collected the samples.

The authors declare that they have no conflict of interest.

## Supplementary Materials

**Suppl Table 1:**
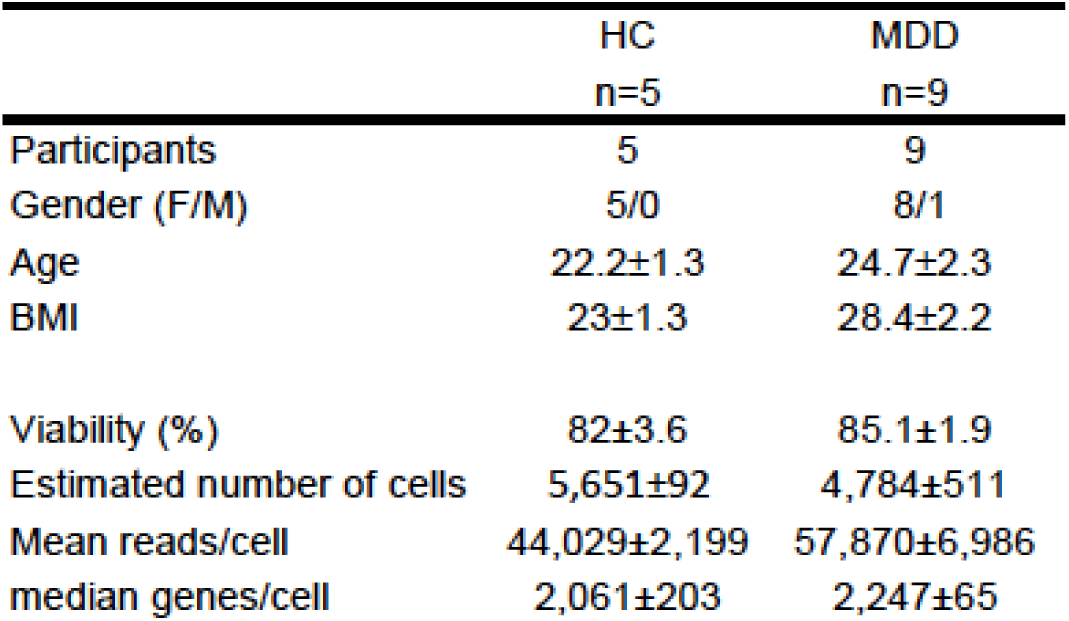
Single-cell RNA sequencing analysis QC found no demographics and cellular confounds.

|  | HC<br>n=5 | MDD<br>n=9 |
| --- | --- | --- |
| Participants | 5 | 9 |
| Gender (F/M) | 5/0 | 8/1 |
| Age | 22.2±1.3 | 24.7±2.3 |
| BMI | 23±1.3 | 28.4±2.2 |
| Viability (%) | 82±3.6 | 85.1±1.9 |
| Estimated number of cells | 5,651±92 | 4,784±511 |
| Mean reads/cell | 44,029±2,199 | 57,870±6,986 |
| median genes/cell | 2,061±203 | 2,247±65 |

**Supplementary Figure 1.**
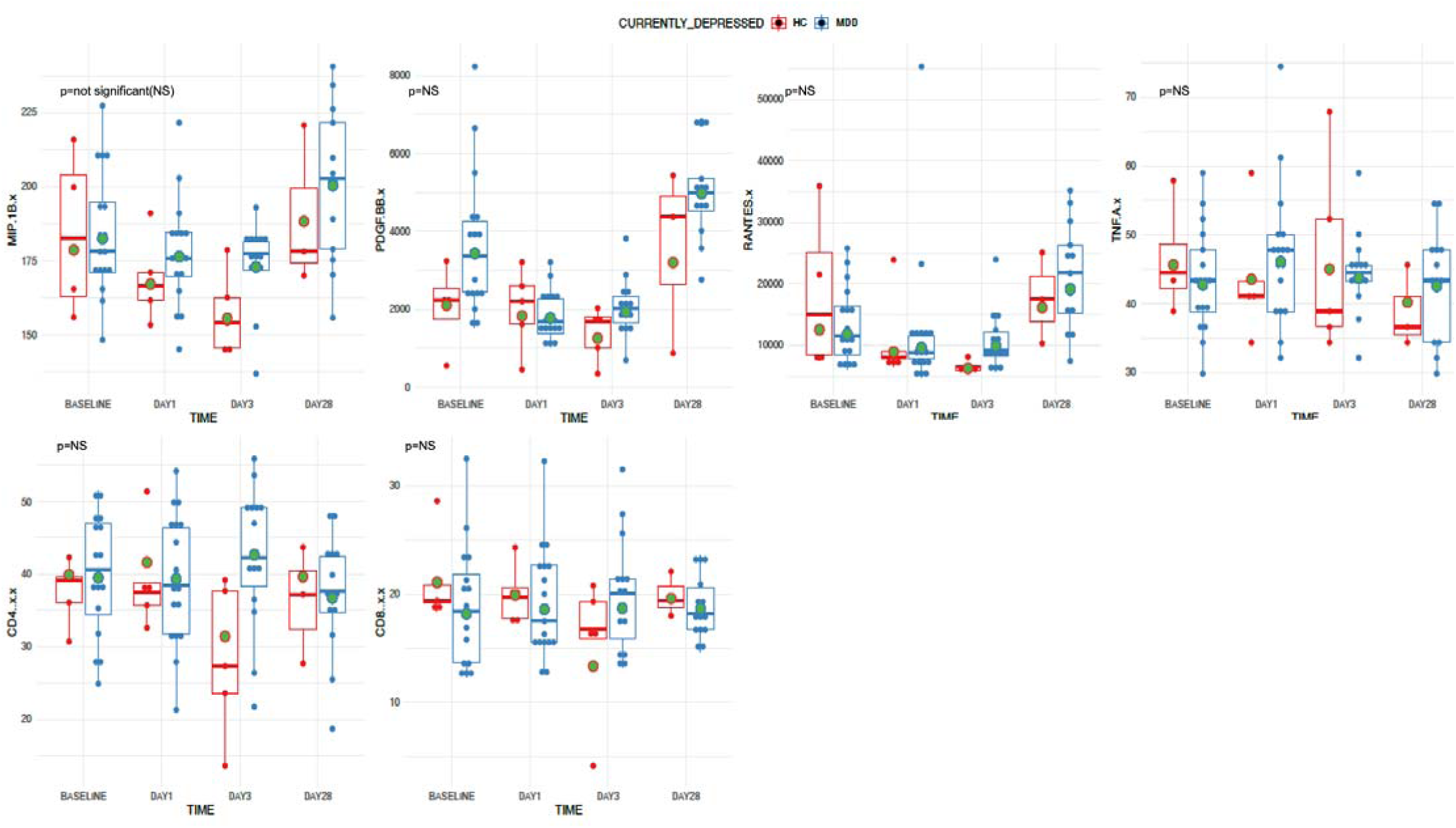
Cytokine, chemokine, and growth factor profiles at baseline and after influenza vaccination in depressed patients and healthy controls. Depressed and healthy control participants donated blood samples before (baseline and after the flu vaccine. The results above show that no group-related differences were observed at baseline or post-flu challenge for all protein-level metrics, including the depicted measures.

## Notes

### Competing Interest Statement

The authors have declared no competing interest.

